# Delineating Medical Education: Bibliometric Research Approach(es)

**DOI:** 10.1101/2021.07.23.453492

**Authors:** Lauren A. Maggio, Anton Ninkov, Jason R. Frank, Joseph A. Costello, Anthony R. Artino

**Affiliations:** Professor of Medicine at Uniformed Services University of the Health Sciences in Bethesda, Maryland, USA @LaurenMaggio; Postdoctoral Fellow at University of Ottawa, School of Information Studies in Ottawa, Ontario Canada @TheNinkov; the Director of Specialty Education for the Royal College of Physicians and Surgeons of Canada, and Vice-Chair Education for the Department of Emergency Medicine, University of Ottawa. @drjfrank; Research Assistant at Uniformed Services University of the Health Sciences in Bethesda, Maryland, USA @jojo_costello; Professor of Health, Human Function, and Rehabilitation Sciences, and Associate Dean for Evaluation and Educational Research at the George Washington University School of Medicine and Health Sciences, Washington, DC, USA @mededdoc

## Abstract

**Background:** The field of medical education remains poorly delineated such that there is no broad consensus of the articles and journals that comprise “the field.” This lack of consensus has implications for conducting bibliometric studies and other research designs (e.g., systematic reviews); it also challenges the field to compare citation scores in the field and across others and for an individual to identify themselves as “a medical education researcher.” Other fields have utilized bibliometric field delineation, which is the assigning of articles or journals to a certain field in an effort to define that field.

**Process:** In this *Research Approach*, three bibliometric field delineation approaches -- information retrieval, core journals, and journal co-citation -- are introduced. For each approach, the authors describe their attempt to apply it in the medical education context and identify related strengths and weaknesses. Based on co-citation, the authors propose the Medical Education Journal List 24 (MEJ-24), as a starting point for delineating medical education and invite the community to collaborate on improving and potentially expanding this list.

**Pearls:** As a research approach, field delineation is complicated, and there is no clear best way to delineate the field of medical education. However, recent advances in information and computer science provide potentially more fruitful approaches to deal with the complexity of the field. When considering these emerging approaches, researchers should consider collaborating with bibliometricians.

Bibliometric approaches rely on available metadata for articles and journals, which necessitates that researchers examine the metadata prior to analysis to understand its strengths and weaknesses, and to assess how this might affect their data interpretation. While using bibliometric approaches for field delineation is valuable, it is important to remember that these techniques are only as good as the research team’s interpretation of the data, which suggests that an expanded research approach is needed to better delineate medical education, an approach that includes active discussion within the medical education community.

## Background

The field of medical education remains poorly delineated. Over the last decade, multiple researchers have aimed to describe medical education and its outputs using bibliometrics, which is the use of statistics to study books, journal articles, and other publication types.^1–6^ To conduct such studies, researchers (including several members of our author team) must make judgment calls about which publications are “in” or “out” of medical education. Researchers who make these calls do so in a fairly ad hoc way because, to our knowledge, there is currently no broad consensus of what constitutes the field of medical education and the articles and journals that comprise it. In this article, we have chosen to refer to medical education as a field, however, we recognize that debate about this distinction exists.^6,7^ This debate, however, is beyond the scope of the present paper.

Our failure to agree upon a common understanding of the field or to delineate -- that is, to portray something precisely or to indicate the exact position, border or boundary^8^ -- the field of medical education has multiple implications not only for conducting bibliometric studies, but also for other research designs like systematic reviews. In addition, our failure to consistently delineate medical education has implications for setting a basis for citation scores and for an investigator’s ability to identify themselves as “a medical education researcher.” Thus, in this manuscript, we describe our attempts to use three *field delineation* approaches: information retrieval, core journals, and journal co-citation. For each approach, we identify strengths and weaknesses and provide practical tips for implementing each approach in medical education. Finally, based on our experiences wrestling with the challenge of field delineation, we invite the medical education community to further collaborate to delineate medical education. To get this conversation started, we introduce a list of 23 journals to serve as a field delineation “starter set” in medical education.

Bibliometric field delineation is described as the assigning of articles or journals to a certain field (i.e., using the field’s “building blocks”^9^ to define that field^10^). Field delineation itself is often viewed as the first step in a research process to allow scientists to explore the structures and dynamics of a research field using bibliometrics.^11,12^ Bibliometrics is the analysis of published information (e.g., journal articles) and its related metadata (e.g, titles, abstracts) using statistics.^13^ Bibliometrics provides a sense of what is valued, recognized and utilized in a field’s scholarly literature.^6^ Several fields, including genomics^14^, nanoscience^11^, and information science^10^ have used field delineation to draw boundaries around their fields and then, using these parameters, have described the field’s journals, topics, members and trends using bibliometrics. To our knowledge, the field of medical education has not been delineated in this systematic way.

There are practical and psychosocial reasons for delineating the field of medical education. For example, if a researcher wishes to conduct a scoping review to identify how a particular theory is used in medical education, they would first need to know the universe of publications within which to look. Similarly, if a researcher wants to understand if medical education is “advancing on big questions” as Regehr^5^ has implored us to do, then they would need to be aware of what is considered “fair game” for inclusion. Without this information it is difficult to chart our progress and build on our previous successes. Field delineation also provides the foundation for generating and understanding benchmark metrics about a field. These metrics (e.g. journal impact factor^15^ or H-index^16^) can be especially important for a field with researchers who may call a variety of academic departments their home. For example, it would be important for a medical educator to clearly communicate to a chair of Medicine, who is responsible for reviewing promotion packets from a broad variety of researchers, the field delineation benchmarks in medical education to demonstrate that their research impact aligns with or surpasses others’ in their specific field.

Field delineation also has several important psychosocial implications for our community. Currently, it can sometimes be unclear who is considered a “member” of the medical education field. This raises issues around whose voices are being heard and whose voices are absent from our ongoing conversations. For example, does medical education have representation from non-English speakers, women, and trainees? Related to this idea of membership and representation, it may be difficult for researchers themselves to claim an identity in medical education, which can confer a sense of belonging and ownership for researchers.^7^

Field delineation is rarely straightforward. Indeed, there is no foolproof approach for all fields and often field borders can be quite fuzzy.^11,12,17^ A field’s border can be especially fuzzy in cases of emerging, interdisciplinary or multidisciplinary fields. In such cases, field delineation can be fraught with additional complications.^10^ For example, in an interdisciplinary field, a given journal may contain articles that address subject matter that cannot be easily assigned to a single field. Furthermore, an article about a given topic, for instance a study of physicians’ social media use, could appear in a medical education journal, a communication journal, or even a general medicine journal. Adding to the complexity of field delineation, it is often the users or actors in a domain who ultimately determine the boundaries of a field, which can introduce additional challenges and biases.^17^

In medical education, we utilize multiple epistemologies and underlying philosophies; lack a specific medical education vocabulary; and our research can and often is very much local to a given educational context. All of these complicating factors make medical education a difficult field to delineate. Nonetheless, we believe it is time to begin making progress toward field delineation in medical education. To that end, in this manuscript we follow the lead of researchers from nanoscience, a similarly multi-disciplinary field, who explored potential field delineation approaches for their field comparing and contrasting the approaches in light of their field’s unique characteristics.^11^ In particular, we describe three approaches and conclude with a proposed “starter set” of medical education journals. In doing so, we intend to prompt future collaborative work to further delineate the field of medical education.

## Process

### Information Retrieval

The information retrieval approach is a popular method of field delineation. For this approach, researchers attempt to identify all of the relevant articles in the field by searching the literature (i.e., information retrieval) such that the retrieved articles would be considered as a representation of the field. We consider this akin to conducting a search as part of a comprehensive systematic review. This approach has been used in several fields (e.g., nanoscience and information science),^9,11,17,18^ as well as in medical education.^3^ In 2010, Lee and colleagues searched PubMed using the medical subject heading (MeSH), “education, medical” as the major focus of articles.

Over a decade later, we loosely replicated Lee et al.’s approach using a broader search approach.^3^ To begin, we conducted a PubMed search for the keywords “medical education.” This search would retrieve any citations with this term in its metadata (e.g., title, abstract, author details), including any articles indexed with the medical subject heading (MeSH) term “education, medical” and its more specific terms related to undergraduate, graduate and continuing medical education. At this point, we considered that this corpus of citations, which contains over 200,000 articles published across hundreds of journals, represents the field of medical education. Notably, we could have chosen to search other databases or multiple databases (e.g., Web of Science, Scopus, Google Scholar), but we chose PubMed for our exploration because it is free, includes MeSH, and is considered the “premier biomedical database”.^19^

While 200,000 citations is a solid seed set of citations, upon closer inspection of the citations retrieved, some limitations were immediately revealed. For example, this search retrieves the article: “Albuminuria intensifies the link between urinary sodium excretion and central pulse pressure in the general population”.^20^ This article, which seems to be unrelated to the field of medical education, is retrieved because the author’s institution is Miyagi University of *Education Medical* Center. We were also concerned about missing relevant articles. For example, *Academic Medicine*, which is often considered a core journal in the field^3,4,6,21^ has published over 12,842 articles since its inclusion in PubMed. However, our strategy only retrieved 5,824 citations from *Academic Medicine* meaning that over 7,000 citations appear to be missing, including the seemingly relevant article: Tolerance for ambiguity among medical students.^22^

After examining the citations, this approach could be optimized by constructing a more comprehensive search string, possibly in collaboration with a medical librarian, to systematically remove some of the irrelevant citations retrieved in the search (e.g., search titles and abstracts only). Researchers could also expand their search by adding additional relevant terms such as “medical student” or “medical school.” Similar to the approach taken in a systematic review, the researchers would most likely need to iterate their search through multiple rounds, which can be a resource-intensive approach.

### Core journals

In a second approach, we identify a collection of “core medical education journals” to define the field. For example, consider the approach undertaken in social work in which the author identified 25 main journals in their field based on Clarivate’s Journal Citation Reports’ (JCR) subject classification.^23^ The JCR classifies over 12,000 journals into subject categories, including the category “social work.” This approach has been considered the “best way” to identify core sets of journals.^24^ However, turning to the JCR, there is no subject category for “medical education” and thus no preset list of journals. There is a somewhat close fit with journals characterized in the category: “education, scientific, disciplines.” However, this also contains titles such as *Engineering Education* and *American Journal of Physics*. Based on the journals’ scope note descriptions and a review of the titles of articles published in 2020, these two journals appear to be outside of the medical education field such that using this approach would introduce a fair amount of irrelevant content. We also investigated a second resource, the Scimago Journal and Country Rank, which includes a seemingly close topic: “social sciences, education.” Similar to the JCR, there were many journals in the resulting list that were well outside the scope of medical education (e.g., *Child Development*).

Next, we considered the “Annotated Bibliography of Journals for Educational Scholarship”, which was collated by the Medical Education Scholarship Research and Evaluation Section (MESRE), a special interest group of the Association of American Medical Colleges.^25^ This list aims to provide researchers and scholars with a sense of the topics, types of manuscripts and the audience for journals in the broad domain of health professions education. The list includes 67 journals and features many of the titles that are commonly referenced in medical education bibliometric studies,^1–3^ which is an encouraging finding. However, it also includes titles that focus on education in general (e.g., *AERA Open)*, allied health disciplines (e.g., *Journal of Dental Education*), and journals that are predominantly clinical, but include some education research (e.g., *JAMA*). While this is an incredibly valuable resource for individuals wanting to identify a place to publish, for our purpose of field delineation, we feel it is too broad. For example, in 2020 *JAMA* published over 15,000 articles of which only 1,886 articles are indexed in PubMed as related to medical education. Therefore, the addition of more than 13,000 seemingly irrelevant articles would introduce quite a bit of noise into the journal set. Additionally, the construction of this bibliography reflects the leanings and preferences of its creators. As an AAMC product and with all the list’s authors based in North America, the list tends to lean heavily towards North American and European publications, with the exception of *Focus on Health Professions Education*, which is the official journal of the Australian and New Zealand Association for Health Professional Educators.

### Journal Co-citation

In a third attempt we utilized journal co-citation. Journal co-citation is the frequency with which two journals are both cited by a third journal. In this case, the two journals both cited by a third journal are considered to be “intellectually related”^26^ (See Figure 1). Co-citation has been defined as a link between two entities (e.g., journals, journal articles, authors) by a third entity citing both.^27^ In other words, co-citation is a measure of the ways in which authors use citations.^26^ For example, in Figure 1 we present an example of a basic case of a co-citation relationship. In paper A, there are citations to both Paper B and Paper C. As a result of these two citations, we would refer to Paper B and Paper C as being “co-cited” by Paper A. This co-citation serves as an indicator that these two papers are likely to be similar to one another. The more instances of Paper B and C being co-cited by other papers (e.g., Paper D, E, and F), the more likely they are to be similar.

**Figure 1.**
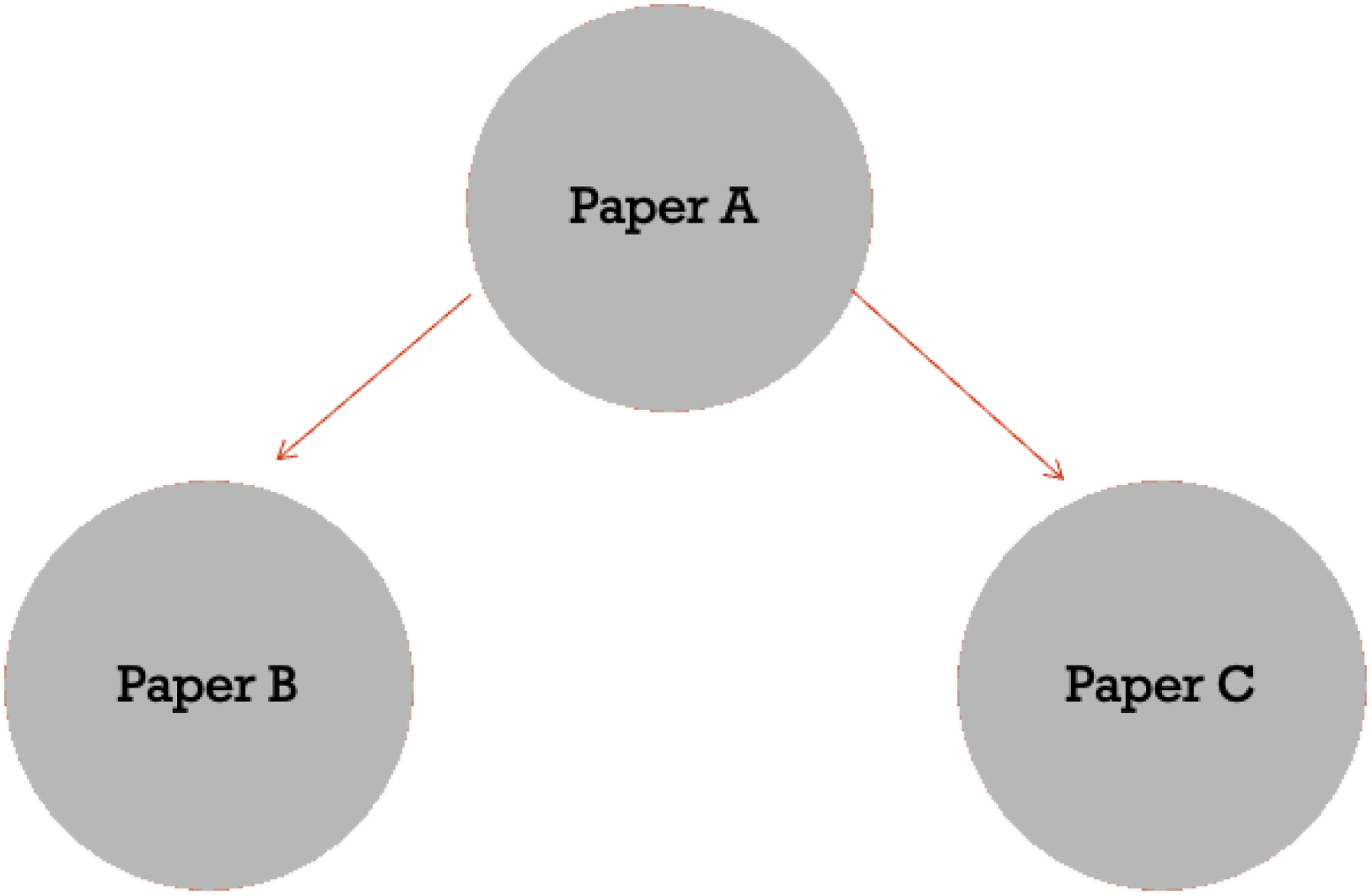
A basic example of co-citation

To conduct co-citation analysis requires a seed set of journals and the metadata of their articles. Since such a set of medical education journals is currently unavailable, we decided to start with the 14 journals that have been described in the literature as “core medical education journals”.^1,3,21^ Using the JCR we determined each journal’s subject categories, which included “education, scientific, disciplines” (n=9), healthcare sciences and services (n=6), education and education research (n=2), medicine, research, experimental (n=1). It is important to note that we focused on the JCR subject classifications to enable an additional step of metadata extraction from Web of Science (WoS). We recognize that this choice introduces limitations that we will discuss later.

For each category, we downloaded the titles of each included journal, which resulted in 987 journals. We screened all journal titles for mentions of education, academia, or teaching in the title. If the title was very generic (e.g., *JAMA*), we reviewed the journal’s scope note to determine if “education” was specifically mentioned. If education was mentioned in the note, the journal was included. This resulted in 24 journals (See online appendix A for journal list). However, at this point our approach hit a roadblock that required a tradeoff. In this set, two of the journals, *Journal of Graduate Medical Education* and the *Canadian Medical Education Journal*, are not indexed in WoS such that we were unable to retrieve the necessary metadata. Thus, these two journals were excluded from the seed set. Additionally, our approach identified the *Journal of General Internal Medicine* (*JGIM*) as being “in scope.” However, between 2000-2020, *JGIM* published 30,783 articles, of which only 2,120 citations included “medical education” when we search all fields. Thus, we made the decision to exclude *JGIM*, since only a minority of its articles (6.8%) focused on medical education. This left 22 journals.

We downloaded from WoS on February 15 and 23, 2021 the metadata for all articles published in these 22 journals between 2000-2020 (n=34,768). Critical to the co-citation approach, the metadata included the references to the articles that had cited the articles published in the 22 journals. To conduct the co-citation analysis we used VOSviewer.^28^ VOSviewer is an open source, freely available software that allows users to construct and visualize bibliometric networks based on co-citation data. This tool has been used in multiple studies.^11,29,30^

Using VOSviewer we identified that there were 66,833 instances of co-citation in our set. Due to the large volume of data, VOSviewer prompted us to select a threshold for displaying co-citations. Thus, we decided to focus on journals with articles that had been co-cited at least 50 times. This resulted in 856 journals, which represented 318,591 citations. For a full listing of journal titles, see deposited data. By the frequency of co-citations the top three journals were *Academic Medicine* (n=44,956), *Medical Education* (n=24,434), and *Medical Teacher* (17,475). These three journals accounted for over 25% of the co-citations (See Table 1). The top 20 journals accounted for over 50% of the co-citations, of which nine journals were from the core set of 22 journals.

**Table 1:**
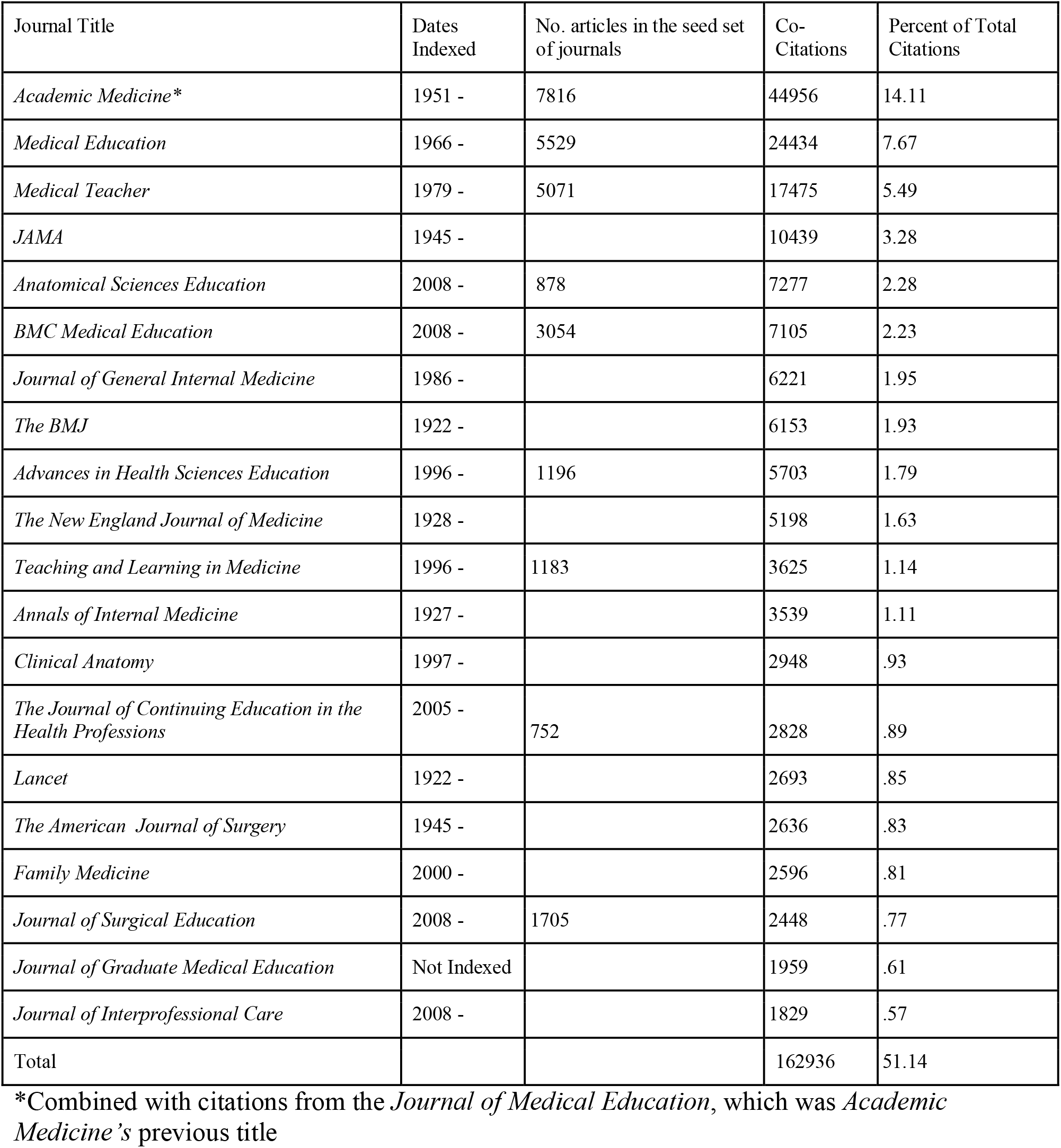
Top 20 journals by number of co-citations. Total co-citations journals = 66,833 with 318,591 citations based on the number of journals that were co-cited at least 50 times.

| Journal Title | Dates Indexed | No. articles in the seed set of journals | Co-Citations | Percent of Total Citations |
| --- | --- | --- | --- | --- |
| *Academic Medicine\** | 1951 - | 7816 | 44956 | 14.11 |
| *Medical Education* | 1966 - | 5529 | 24434 | 7.67 |
| *Medical Teacher* | 1979 - | 5071 | 17475 | 5.49 |
| *JAMA* | 1945 - |  | 10439 | 3.28 |
| *Anatomical Sciences Education* | 2008 - | 878 | 7277 | 2.28 |
| *BMC Medical Education* | 2008 - | 3054 | 7105 | 2.23 |
| *Journal of General Internal Medicine* | 1986 - |  | 6221 | 1.95 |
| *The BMJ* | 1922 - |  | 6153 | 1.93 |
| *Advances in Health Sciences Education* | 1996 - | 1196 | 5703 | 1.79 |
| *The New England Journal of Medicine* | 1928 - |  | 5198 | 1.63 |
| *Teaching and Learning in Medicine* | 1996 - | 1183 | 3625 | 1.14 |
| *Annals of Internal Medicine* | 1927 - |  | 3539 | 1.11 |
| *Clinical Anatomy* | 1997 - |  | 2948 | .93 |
| *The Journal of Continuing Education in the Health Professions* | 2005 - | 752 | 2828 | .89 |
| *Lancet* | 1922 - |  | 2693 | .85 |
| *The American Journal of Surgery* | 1945 - |  | 2636 | .83 |
| *Family Medicine* | 2000 - |  | 2596 | .81 |
| *Journal of Surgical Education* | 2008 - | 1705 | 2448 | .77 |
| *Journal of Graduate Medical Education* | Not Indexed |  | 1959 | .61 |
| *Journal of Interprofessional Care* | 2008 - |  | 1829 | .57 |
| Total |  |  | 162936 | 51.14 |

\*Combined with citations from the *Journal of Medical Education*, which was *Academic Medicine's* previous title

The 22 journals from the initial set accounted for 41.2% of co-citations. Although, due to database constraints noted above, we excluded two journals, *JGME* and the *Canadian Journal of Medical Education,* both were identified in the literature as “core journals”;^3,21^ these journals were co-cited and we have included them in Table 2.

**Table 2:** Representation of the MEJ-24, which comprises journals in the core set of 22 medical education journals plus the *Journal of Graduate Medical Education* and *Canadian Medical Education Journal*.

| Rank | Journal | No. articles in the seed set of journals | Co-citations | Percent of Total |
| --- | --- | --- | --- | --- |
| 1 | *Academic Medicine* | 7816 | 44956 | 14.11 |
| 2 | *Medical Education* | 5529 | 24434 | 7.67 |
| 3 | *Medical Teacher* | 5071 | 17475 | 5.49 |
| 5 | *Anatomical Sciences Education* | 878 | 7277 | 2.28 |
| 6 | *BMC Medical Education* | 3054 | 7105 | 2.23 |
| 9 | *Advances in Health Sciences Education* | 1196 | 5703 | 1.79 |
| 11 | *Teaching and Learning in Medicine* | 1183 | 3625 | 1.14 |
| 14 | *Journal of Continuing Education in the Health Professions* | 752 | 2828 | .89 |
| 18 | *Journal of Surgical Education* | 1705 | 2448 | .77 |
| 19 | *Journal of Graduate Medical Education* | 0 | 1959 | .61 |
| 21 | *Clinical Teacher* | 1803 | 1819 | .57 |
| 29 | *Medical Education Online* | 600 | 1343 | .42 |
| 37 | *GMS Journal for Medical Education* | 394 | 1207 | .38 |
| 45 | *Simulation in Healthcare* | 781 | 961 | .30 |
| 48 | *Advances in Medical Education and Practice* | 981 | 928 | .29 |
| 52 | *Education for Health* | 548 | 815 | .26 |
| 60 | *Perspectives on Medical Education* | 612 | 686 | .22 |
| 67 | *International Journal of Medical Education* | 485 | 629 | .20 |
| 125 | *Journal of Educational Evaluation for Health Professions* | 372 | 346 | .11 |
| 211 | *African Journal of Health Professions Education* | 392 | 199 | .06 |
| 395 | *Journal of Medical Education and Curricular Development* | 272 | 105 | .03 |
| 454 | *Canadian Medical Education Journal* | 0 | 91 | .03 |
| 630 | *Focus on Health Professional Education* | 152 | 68 | .02 |
| 677 | *BMJ Simulation & Technology Enhanced Learning* | 192 | 62 | .02 |
|  | Total | 34768 | 133290 | 41.84 |

\*This table does not include the *Journal of General Internal Medicine*

Despite a lot of effort this co-citation approach also has limitations. First the seed set of journals excluded journals not indexed in the JCR, including *JGME* and the *Canadian Medical Education Journal*. However, as both of these journals were both identified in the literature as core journals in medical education^3,21^ and together accounted for .64% of the total co-citations, we feel both of these publications warrant inclusion in the field of medical education. Additionally, due to indexing limitations, this approach does not take into account most specialty journals that focus on education, which tend to be indexed in relation to their speciality only. For example, *Academic Pediatrics* is indexed in only the category of Pediatrics despite the fact that the journal’s scope note describes it as an “active forum for the presentation of pediatric educational research”.^31^

From a methods standpoint, a benefit of co-citation is that it provides several ways of thinking about defining a field’s core set of journals; however, this is also a limitation in that there is no gold standard approach to determining how to best interpret the results. In this article, we provided two interpretations, which did not produce what we would consider the “ideal set of journals” to define the field of medical education. The first interpretation is based on the “top 20 journals”. This set includes nine journals from the core set, but also introduces clinical journals (e.g., *JAMA, BMJ*). Although these clinical journals have some coverage of medical education, medical education research is a minority of the content covered in these journals. Thus, by including these clinical journals, we also introduce a good deal of irrelevant content.

The second interpretation was an attempt to determine how the original core set of journals performed. In other words, we tried to determine if these 22 journals greatly contributed to co-citations. Because these journals plus *JGME* and the *Canadian Journal of Medical Education* contributed to 41.84% of the co-citations, we propose that while not an “ideal set” of journals, that they represent a starting point for delineating the field. We call this journal set the Medical Education Journals-24 (MEJ-24), based on the year it was first described.

## Pearls

Notwithstanding our best efforts, we agree with Munoz that there is no perfect means of field delineation^11^ – at least not for a field like medical education with several complicating factors. In Table 3, we provide a listing of the pros and cons for each of the three approaches we attempted in this paper. Next, while we have tried to embed “practical pearls of wisdom” throughout the manuscript, we focus on several key considerations for those considering similar projects using bibliometrics approaches and for those seeking to broadly delineate the field of medical education.

**Table 3:** Summary table of approaches described for delineating the field of medical education and the related pros and cons.

| Approach | Pros | Cons |
| --- | --- | --- |
| Information Retrieval | Retrieves citations from across a range of journals<br>Low-ish effort<br>From citation data a researcher could begin to characterize the field (author characteristics, publication types, topics, etc) | Retrieves papers that are outside of scope<br>Misses relevant papers<br>Generally requires searching multiple databases, some of which require a subscription |
| Definitive Journals | Retrieves citations only from the included journals<br>A researcher could mine the journal set for publications that could be used to characterize the field<br>If a definitive journal set exists this is low effort | Exclude articles not published in the journal set<br>Contains the bias of those that created the journal set<br>It is difficult to balance journals that include some medical education articles vs. those that focus on medical education |
| Co-citation | Retrieves citations from across a range of journals<br>Highly used and validated<sup>26</sup><br>Offers a method to identify most important journals<sup>26</sup> | A high-effort approach that may require consultation with a bibliometrician<br>May require access to subscription databases<br>Can miss journals that are related, but that do not cite the same resource<br>Only works for articles with references and citations, which makes this less efficacious with newer articles that have not accrued citations |

The three approaches described here have been used for field delineation for many years. However, recent advances in information and computer science have enabled researchers to expand these approaches. For example, researchers have used social network analysis and natural language processing to help make sense of the increasing amounts of available data.^12^ To this end, we encourage those interested in field delineation to explore emerging methods with the caveat that they strongly consider collaborating with researchers with expertise in information science, specifically those with expertise in bibliometrics.

In each approach, we necessarily relied on the available metadata for articles and journals. As we observed, this can be problematic for all three approaches. Therefore, it is important for researchers to examine the metadata prior to analysis to understand the strengths and weaknesses of their data set and to assess how this might impact their interpretations of the data. Furthermore, we would encourage journal editors to investigate how their journal is indexed. For example, should the editor of the *Journal of Academic Pediatrics*, which describes an education mission, seek to be indexed in Web of Science as an education journal in addition to its current indexing as only pediatrics? In addition to facilitating field delineation research, this may also facilitate the findability of the journal’s content by those using educational search terms.

While the use of bibliometric approaches for field delineation are valuable it is important to bear in mind that these techniques are only as good as the research team’s interpretation of that data. Therefore, we believe an expanded research approach is needed, one that includes active discussion between a wide diversity of medical education stakeholders. We recommend that this discussion be structured with the aim of arriving at a working consensus on the scope of the field. To this end, we call the community to action to use the MEJ-24 as a starting point or seed set of journals to inform this critical conversation.

## Ethical Approval

Reported as not applicable

## Data

None reported

## Disclaimer

The views expressed in this article are those of the authors and do not necessarily reflect the official policy or position of the Uniformed Services University of the Health Sciences, the Department of Defense, or the U.S. Government.

## References

1. Maggio LA, Costello JA, Norton C, Driessen EW, Artino AR Jr. Knowledge syntheses in medical education: A bibliometric analysis. Perspect Med Educ. 2021;10(2):79–87. doi:10.1007/s40037-020-00626-9

2. Rotgans JI. The themes, institutions, and people of medical education research 1988-2010: content analysis of abstracts from six journals. Adv Health Sci Educ Theory Pract. 2012;17(4):515–527. doi:10.1007/s10459-011-9328-x

3. Lee K, Whelan JS, Tannery NH, Kanter SL, Peters AS. 50 years of publication in the field of medical education. Med Teach. 2013;35(7):591–598. doi:10.3109/0142159X.2013.786168

4. Azer SA. The top-cited articles in medical education: a bibliometric analysis. Acad Med. 2015;90(8):1147–1161. doi:10.1097/ACM.0000000000000780.

5. Regehr G. Trends in medical education research. Acad Med. 2004;79(10):939–947. doi:10.1097/00001888-200410000-00008

6. Albert M, Rowland P, Friesen F, Laberge S. Interdisciplinarity in medical education research: myth and reality. Adv Health Sci Educ Theory Pract. 2020;25(5):1243–1253. doi:10.1007/s10459-020-09977-8

7. ten Cate, O. Health professions education scholarship: The emergence, current status, and future of a discipline in its own right. FASEB BioAdvances. 2021;3:510–522. doi:10.1096/fba.2021-00011

8. Delineate. The Oxford English Dictionary. Oxford Dictionaries. https://www.google.com/search?q=define+delineate Retrieved July 19, 2021.

9. Zitt, M. Meso-level retrieval: IR-bibliometrics interplay and hybrid citation-words methods in scientific fields delineation. Scientometrics. 2015;102:2223–2245. doi: 10.1007/s11192-014-1482-5

10. Zhao D. Mapping library and information science: Does field delineation matter? Proceedings of the American Society for Information Science and Technology. 2009;46(1):1–11. doi: 10.1002/meet.2009.1450460279

11. Muñoz-Écija T, Vargas-Quesada B, Rodríguez ZC. Coping with methods for delineating emerging fields: Nanoscience and nanotechnology as a case study. Journal of Informetrics. 2019;13(4):100976. doi:10.1016/j.joi.2019.100976.

12. Lietz H. Drawing impossible boundaries: field delineation of Social Network Science. Scientometrics. 2020;125(3):2841–76. doi:10.1007/s11192-020-03527-0

13. Broadus RN. Toward a definition of “bibliometrics”. Scientometrics. 1987;12(5-6):373–379. doi: 10.1007/BF02016680

14. Laurens P, Zitt M, Bassecoulard E. Delineation of the genomics field by hybrid citation-lexical methods: interaction with experts and validation process. Scientometrics. 2010;82:647–662. doi: 10.1007/s11192-010-0177-9

15. Garfield E. The history and meaning of the journal impact factor. JAMA. 2006;295(1):90–93. doi:10.1001/jama.295.1.90

16. Hirsch JE. An index to quantify an individual’s scientific research output. Proc Natl Acad Sci USA. 2005;102(46):16569–16572. doi:10.1073/pnas.0507655102

17. Glänzel W. Bibliometrics-aided retrieval: where information retrieval meets scientometrics. Scientometrics. 2015;102:2215–2222. doi: 10.1007/s11192-014-1480-7

18. Aksnes DW, Olsen TB, Seglen PO. Validation of Bibliometric Indicators in the Field of Microbiology: A Norwegian Case Study. Scientometrics. 2000;49(1):7–22. doi: 10.1023/A:1005653006993

19. Masic I, Milinovic K. On-line biomedical databases-the best source for quick search of the scientific information in the biomedicine. Acta Inform Med. 2012;20(2):72–84. doi:10.5455/aim.2012.20.72-84

20. Tagawa K, Tsuru Y, Yokoi K, Aonuma T, Hashimoto J. Albuminuria intensifies the link between urinary sodium excretion and central pulse pressure in the general population: the Wakuya study [published online ahead of print, 2021 Apr 24]. Am J Hypertens. 2021;hpab057. doi:10.1093/ajh/hpab057

21. Maggio LA, Leroux TC, Meyer HS, Artino AR Jr. #MedEd: exploring the relationship between altmetrics and traditional measures of dissemination in health professions education. Perspect Med Educ. 2018;7(4):239–247. doi:10.1007/s40037-018-0438-5

22. Geller G, Faden RR, Levine DM. Tolerance for ambiguity among medical students: implications for their selection, training and practice. Soc Sci Med. 1990;31(5):619–624. doi:10.1016/0277-9536(90)90098-d

23. Martínez MA, Cobo MJ, Herrera M, Herrera-Viedma E. Analyzing the Scientific Evolution of Social Work Using Science Mapping. Research on Social Work Practice. 2015;25(2):257–277. doi:10.1177/1049731514522101

24. Pudovkin AI, Garfield E. Algorithmic procedure for finding semantically related journals. Journal of the American Society for Information Science and Technology. 2002;53(13):1113–1119. doi:10.1002/asi.10153

25. Association of American Medical Colleges, Regional Groups on Educational Affairs (GEA): Medical Education Scholarship, Research and Evaluation Section. Annotated Bibliography of Journals for Educational Scholarship, Berry, ed. https://www.aamc.org/media/38166/download. Revised 2019. Accessed July 8, 2021.

26. Zupic I, Cater T. Bibliometric methods in management and organization. Organizational Research Methods. 2015;18(3):429–472.

27. Small H. Co-citation in the scientific literature: A new measure of the relationship between two documents. Journal of the American Society for Information Science. 1973;24(4):265–269. doi: 10.1002/asi.4630240406

28. van Eck NJ, Waltman L. Software survey: VOSviewer, a computer program for bibliometric mapping. Scientometrics. 2010;84(2):523–538.

29. Hallinger P, Kovačević J. A Bibliometric Review of Research on Educational Administration: Science Mapping the Literature, 1960 to 2018. Review of Educational Research. 2019;89(3):335–369. doi:10.3102/0034654319830380.

30. Flis I, van Eck NJ. Framing psychology as a discipline (1950-1999): A large-scale term co-occurrence analysis of scientific literature in psychology. Hist Psychol. 2018;21(4):334–362. doi:10.1037/hop0000067

31. Academic Pediatric Association. Academic Pediatrics: Aims and Scope. https://www.academicpedsjnl.net/content/aims. Accessed July 8,2021.

